# DEBrowser: Interactive Differential Expression Analysis and Visualization Tool for Count Data

**DOI:** 10.1101/399931

**Authors:** Alper Kucukural, Onur Yukselen, Deniz M Ozata, Melissa J Moore, Manuel Garber

## Abstract

**Background:** Sequencing data has become a standard measure for studying diverse cellular activities. For example, gene expression is accurately measured by RNA sequencing (RNA-Seq) libraries, protein-DNA interactions are captured by chromatin immunoprecipitation sequencing (ChIP-Seq), protein-RNA interactions by crosslinking immunoprecipitation (CLIP-Seq) or RNA immunoprecipitation (RIP-Seq) sequencing, DNA accessibility by assay for transposase-accessible chromatin (ATAC-Seq), and DNase or MNase sequencing libraries. Analysis of these sequencing techniques involve library-specific approaches. However, in all cases, once the sequencing libraries are processed, the result is a count table specifying the estimated number of reads originating from a genomic locus. Differential analysis to determine which loci have different cellular activity under different conditions starts with the count table and iterates through a cycle of data assessment, preparation and analysis. Such iterative approach relies on multiple programs and is therefore a challenge for those without programming skills.

**Results:** We developed DEBrowser, as an R bioconductor project, to interactively visualize each step of the differential analysis of count data, without any requirement for programming expertise. The application presents a rich and interactive web based graphical user interface based on R’s shiny infrastructure. We use shiny’s reactive programming interface for a dynamic webpage that responds to user input and integrates its visualization widgets at each stage of the analysis. In this way, every step of the analysis can be displayed in one application that combines many approaches and multiple results. We show DEBrowser’s capabilities by reproducing the analysis of two previously published data sets.

**Conclusions:** DEBrowser is a flexible, intuitive, web-based analysis platform that enables an iterative and interactive analysis of count data without any requirement of programming knowledge.

## Background

Sequencing techniques have been widely used to measure the activity of genomic regions across conditions. Typical uses include differential expression [1–3], small RNA abundances [4,5], epigenetic state [6,7], protein/RNA interactions [8–10] and DNA/RNA interactions [11,12]. Even though each of these sequencing libraries requires very specific processing steps to determine the genomic loci underlying the observed sequencing reads [13–15], the outputs are always count matrices with rows representing the genomic features of interest such as genes, exons, DNA accessible or DNA bound regions, and the columns being the samples. The values in this table are the number of mapping reads to each defined locus for each sample. The most common first step in analysis of such tables is “Which loci exhibit significant differences between different groups of samples?”. Differential analysis of count data typically involves an iterative approach that heavily relies on visualization and unsupervised statistical analysis. A typical analysis consists of an iterative application of three main tasks; data assessment, data preparation, and differential expression analysis (Fig 1).

**Fig. 1.**
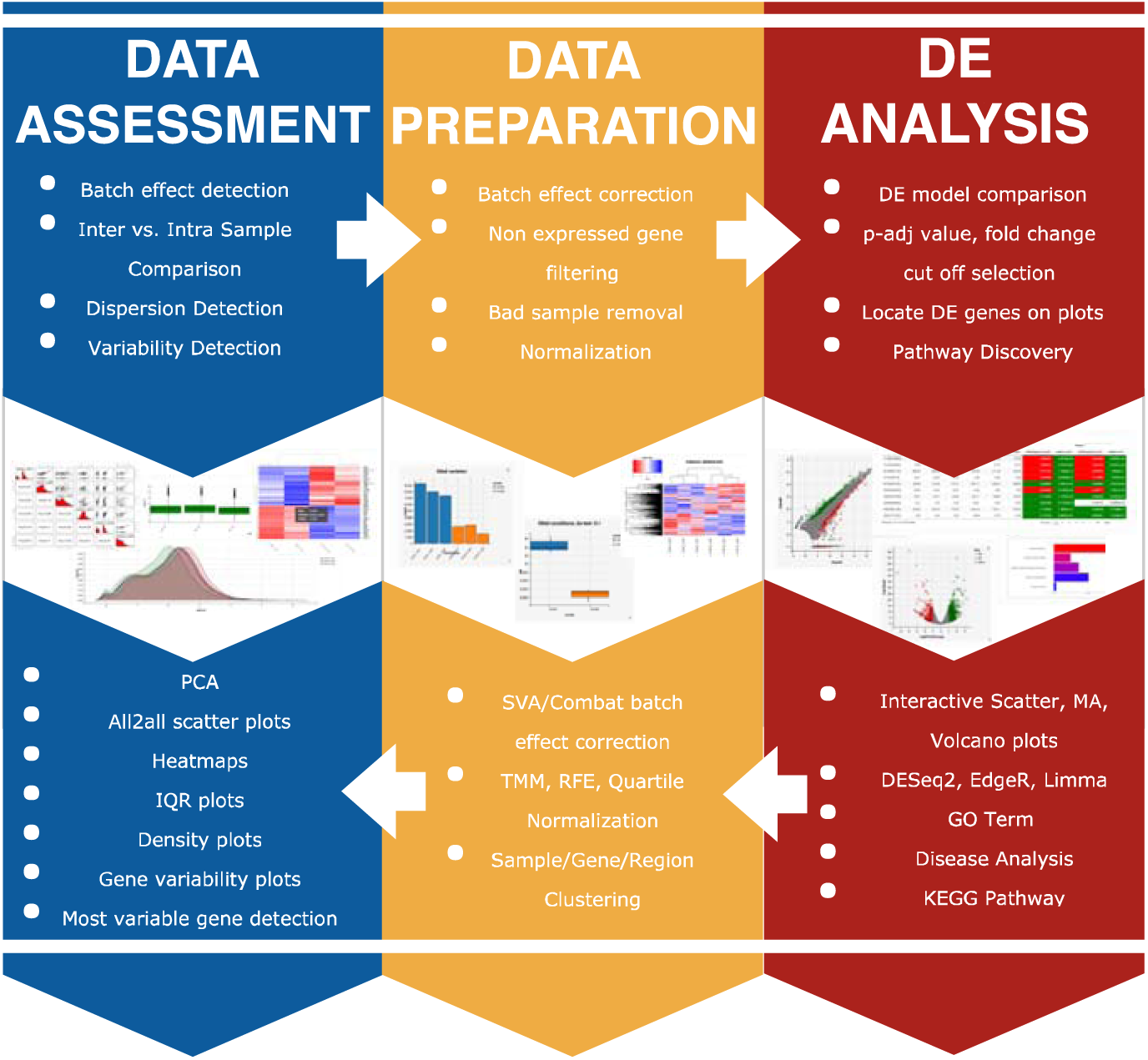
With DEBrowser, an analysis’ stages are presented in one package.

**Data assessment** evaluates the impact of latent factors that may not be related to biological differences. Such differences might come from technical factors, such as DNA/RNA fragmentation, the number of PCR cycles, sequencing depth, these may altogether confound the actual biologically relevant differences presenting in samples [16,17]. Therefore, the count matrix represents a combination of both the biological and technical variation. Unsupervised methods such as clustering and principal component analysis (PCA) are powerful ways to identify technical sources of variance [18–20]. Data assessment involves the use of standard unsupervised tools for inter-vs. intra-sample comparisons to assess the impact of sources of unwanted variability, such as batch effects, low quality samples, and samples with low sequencing depth. **Data preparation** builds on data assessment to determine and then apply the best approaches to reduce the impact of unwanted sources of variance. Typically that includes the elimination of low quality samples, and pre-filtering - removal of features having low counts, that increase the variability resulting from non-biological sources [21–25]. Data preparation may also include batch effect correction [26], which removes the sources of variation in the groups of samples due to technical differences. Following these steps leads to more accurate differential analysis.

We created DEBrowser to enable iterative analysis by non-programmers to achieve similar results to those obtained by investigators well versed in R programming language. DEBrowser facilitates a modular iterative analysis and visualization process through an intuitive user interface. It integrates multiple algorithms and visualization techniques. The goal is to allow the users to iteratively inspect and apply each of the many approaches comprised of the three stages described above. DEBrowser provides an evaluation of the results at each analysis step, and determines whether further improvements are necessary. DEBrowser goes further than providing non-interactive plots or heatmaps: it allows users to explore any anomaly or potential result by zooming-in on data subsets, to select or hover over any regions or genes and to plot the heatmap, bar or box plots to visualize the sample variation interactively. For example, users may select the most differentially expressed genes on a volcano plot, and re-display them in a heatmap, that can be further used to inspect the expression of each individual genes across all replicates.

### DE Analysis and Visualization packages

A number of graphical user interfaces address the need for user-friendly, programming-free visualizations [27–33] (Table S1). However, all of these approaches have limited interactivity for users to carry out more sophisticated analysis. Similar to DEBrowser, existing tools accept count data as input to visualize, identify, and perform differential analysis and gene ontology. They also visualize results using scatter, MA or volcano plots, as well as heatmaps and PCA plots. DEBrowser goes one step further by enabling hands-on manipulation of the data and by enabling users to re-plot a selected subset of data. These features make DEBrowser a sophisticated tool for data exploration. DEBrowser allows users to color the genes that exceed the different fold change cutoffs after differential expression analysis with only a few clicks. Furthermore, all plots are immediately redrawn upon detection of changes in the plotting parameters or after any data subsetting operation. Similarly, for easy access to the underlying data, DEBrowser supports hovering to obtain detailed information of individual data points.

In contrast to existing visualization and differential analysis tools, DEBrowser focuses on data assessment and preparation: it intrinsically supports normalization and batch correction methods. Once the users identify a clear bias in data preparation, processing or sequencing, DEBrowser allows the users to minimize the differences unrelated to the actual biological differences by the experiment [26,34–38]. This capability is intended to address the needs of large projects that process samples over an extended period of time, and to help users to compare samples available from public repositories originating from different laboratories.

## Implementation

DEBrowser is implemented in R as a shiny application using generic shiny components and layouts [39]. In particular, DEBrowser relies on R’s plotly package [40] both for interactive plots and to display multi-panel data. For heatmaps, we used heatmaply package [41].

DEBrowser relies on shiny’s reactive programming model to detect changes in any input control, plot, or bound object so that users can interactively explore their results and investigate the thresholds. When a change in a plot parameter is detected, plots bound to that input are notified and redrawn. As a result, with few clicks, users can change the highlighted genes from the DE results that exceed a significance of 0.05 and a 2-fold change, to those exceeding a significance of 0.01 and a 10-fold change.

### Design and Key Features

To show the general applicability of DEBrowser on “count data” from different data types; we used a large data set that we recently generated to study gene regulation in innate immune cells (human monocyte derived dendritic cells, hMDDCs) in response to Toll-like receptor signaling [42]. This study generated RNA-Seq, ChIP-Seq and ATAC-Seq [43] to track changes in transcription and regulatory element activity in the course of Toll-like receptor (TLR) signaling.

These data are ideal to showcase the main features of DEBrowser and how DEBrowser can be used throughout the analysis cycle. Indeed we show how DEBrowser was used for data assessment to identify batch effects, and data preparation by filtering low count features and removing batch effects.

#### A. Data assessment

**Quality control (QC):** The quality control of the count data is a fundamental step in analysis, yet it is not well supported in current applications. Users can easily use DEBrowser to establish the suitability of the data for differential analysis and to determine whether further normalization, batch correction or sample removal are necessary. To this end, DEBrowser implements PCA, all2all scatter, heatmaps, interquartile range (IQR) and density plots. These plots can be drawn using a user defined subset of genes, for example by choosing the top N variable genes [40,41]. The subset of genes can be defined graphically by either and expression cutoff or by directly selecting them from another plot.

For example, the data for hMDDCs show clear donor dependent differences, which are visible in all2all plots and are captured by the second principal component (Fig. 2a). These differences may be the result of genetic heterogeneity or simply due to technical variability in library construction. Regardless of the source, the variability introduced by inter-donor differences have a direct impact on detecting TLR responsive genes.

**Fig. 2.**
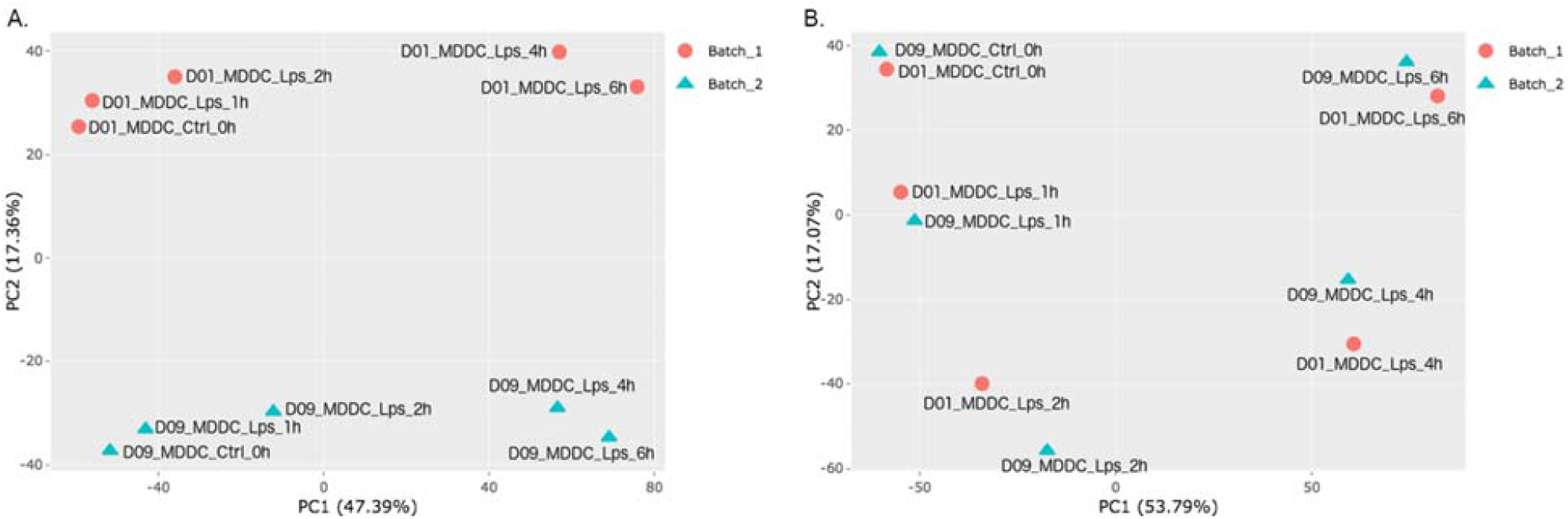
PCA plot (**a**) before and (**b**) after batch effect correction. Batch 1 and 2 are represented by circles and triangles, respectively.

#### Plots available for data assessment

**Principal Component Analysis (PCA):** PCA finds an ordered coordinate transformation whose bases capture, in decreasing order, the most variance in the data. DEBrowser allows users to plot any pair of principal components in a scatter plot. Once users specify sample information (e.g. condition), DEBrowser uses colors or shapes to group samples. These plots are ideal to detect outliers or batch effects (Fig. 2).

**All2all scatter plots:** Gives an overview of sample similarities and variance by plotting all pairwise scatter plots (Fig. S1). Low correlation or high variance across replicates will negatively impact the power to detect DE.

**Heatmaps:** DEBrowser lets users select genes based on variance, minimum expression level, DE p-value, or after manually selecting a set of genes from any gene centric plots (e.g. scatter, volcano, and other heatmaps). Heatmaps are also useful to assess replicate variability, low quality samples, or batch effects (Fig. 3). Similar to other plots, heatmaps can be used to visualize any type of count data and are ideal to identify global patterns in the data such as dynamic changes in chromatin accessibility following TLR signaling [42] (Fig. S2).

**Fig. 3.**
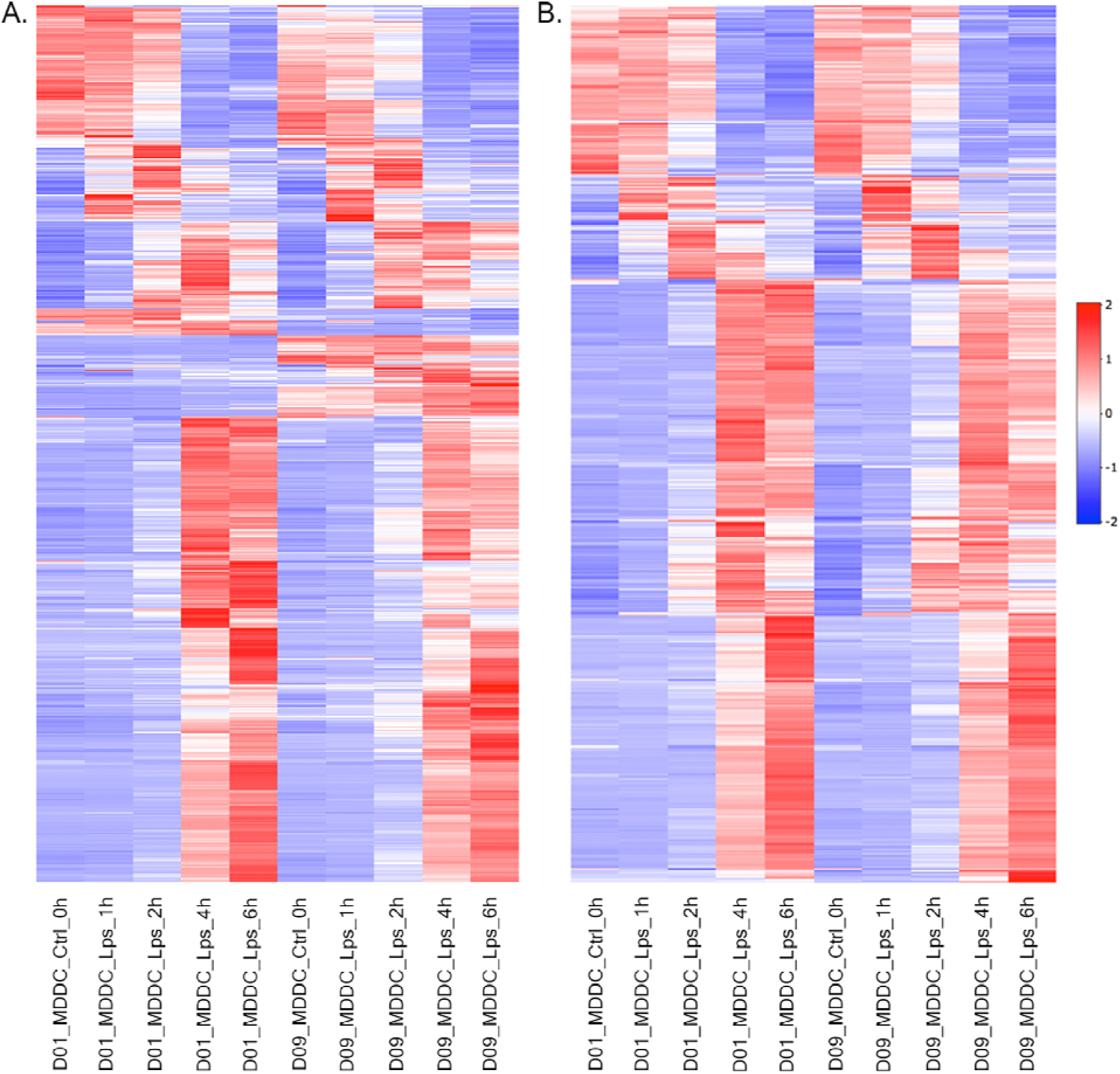
Heatmap for RNA-Seq data at different time points from two donors before (**a**) and after (**b**) batch correction. Heatmaps show the top 1000 most varied genes (based on coefficient of variance) whose total counts are higher than 100.

**IQR and Density plots:** Interquartile range and density plots display a sample’s quantification distribution in two different ways. Using these plots, users can detect any global discrepancy across samples and evaluate the impact of normalization on the distribution of counts. DEBrowser simplifies comparisons by providing plots for both normalized and unnormalized data. Plots are re-drawn as soon as users change the normalization method.

#### B. Data preparation

**Removal of low expression features:** Removing features (genes, or genomic regions) that have low coverage due for example to their low expression, increases the speed and accuracy of DE algorithms. It also helps perform more accurate dispersion calculations and multiple hypothesis correction [44]. DEBrowser provides three common ways to filter these features: by specifying a minimum signal in at least one sample, by a minimum average signal across all samples or by requiring a minimum signal in at least n samples (n defined by the users). Once a filtering criteria is specified, DEBrowser reports read counts for each sample (Fig. 4a,b) and plots the feature count distributions before and after applying the filtering (Fig. 4c,d).

**Fig. 4.**
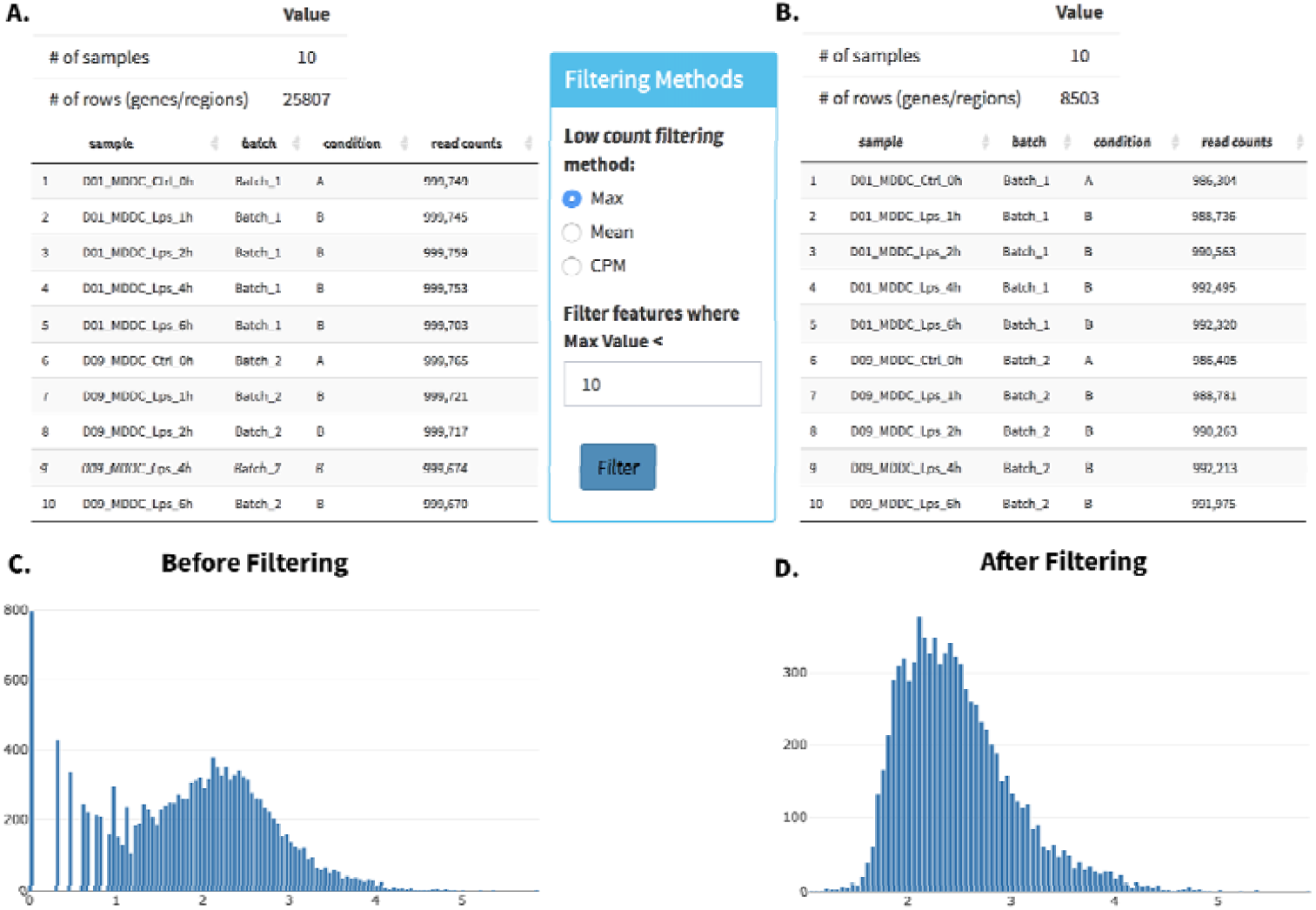
Comparing the number of genes, read counts (a, b) and their distributions (c, d) before and after filtering.

**Normalization:** The count data originating from a sequencing experiment is affected by sequencing depth as well as from differences in the composition of the detected features [45,46]. DEBrowser supports normalization methods specifically designed for count data: median ratio normalization (MRN) implemented in DESeq2 package [44,47], Trimmed Mean of M-values (TMM), Relative Log Expression (RLE), and upper quartile methods implemented in the EdgeR package [48]. To evaluate the effect of normalization, DEBrowser immediately displays PCA, IQR and density plots after normalization.

**Batch effect correction:** When quality control shows a clear batch effect that can be traced back to a technical artifact (e.g. different sequencing devices, different personnel, library kits, reagent batch) DEBrowser allows the users to minimize the batch effect, whenever the experimental design allows batch correction [49]. DEBrowser allows users to specify a batch for every sample via a simple tab separated file that can be created using a text editor or spreadsheet software. Given a batch specification, DEBrowser supports two different methods: ComBat [26,50] and Harman [51] to remove nuisance sources of variance. The resulting, batch corrected dataset can be evaluated using the same tools available for initial quality control. The detected batch effect in Fig. 2b is corrected using batch effect correction module after DESeq2 normalization. It clearly shows how effective was the batch effect correction is if they both share a common sample that are supposed to be in the same state. In this case the control samples are not treated and after the correction, we observe all the corresponding time points cluster together in PCA plot in Fig. 2b. In DEBrowser, all of these steps can be visualized to evaluate the result of any batch correction or normalization method applied.

#### C. Differential expression analysis

To demonstrate a typical usage of DEBrowser, we applied DEBrowser on a data set from a previously published study on the role of Jun terminal kinases (JNK1 and JNK2) in the liver and their role in insulin resistance [2]. For this purpose the authors relied on four different mouse genotypes: wild type (WT), and hepatocyte specific knockouts of Jnk1 and Jnk2 independently (L^Δ 1^, L^Δ2^), and double knockout (L ^Δ1,2^). Each genotype was fed either a regular or a high fat diet. Thereafter, hepatic expression was assayed for each genotype fed with corresponding diet in triplicate using RNA-Seq, resulting in a total of 24 libraries. This study is ideal for DE analysis as it included three replicates per condition. Therefore, we used RSEM [15] for library quantification and DEBrowser to analyze the resulting read count table.

DEBrowser supports differential analysis of count data using DESeq2 [44], EdgeR [48], and Limma [52]. Users can perform differential analysis by defining the groups of samples to compare. DE results can be visualized through the same scatter, volcano and MA plots used for data assessment (Fig. 5a-c). Users can highlight results by specifying desired significance and fold change cut-offs. All plots allow interactive access: Users may select a point within the plot to zoom-in and re-display only selected data. Plots are redrawn as soon as the users change any parameter or selects points to zoom-in on any data point or sets of points that can be investigated by graphically selecting them.

**Fig. 5.**
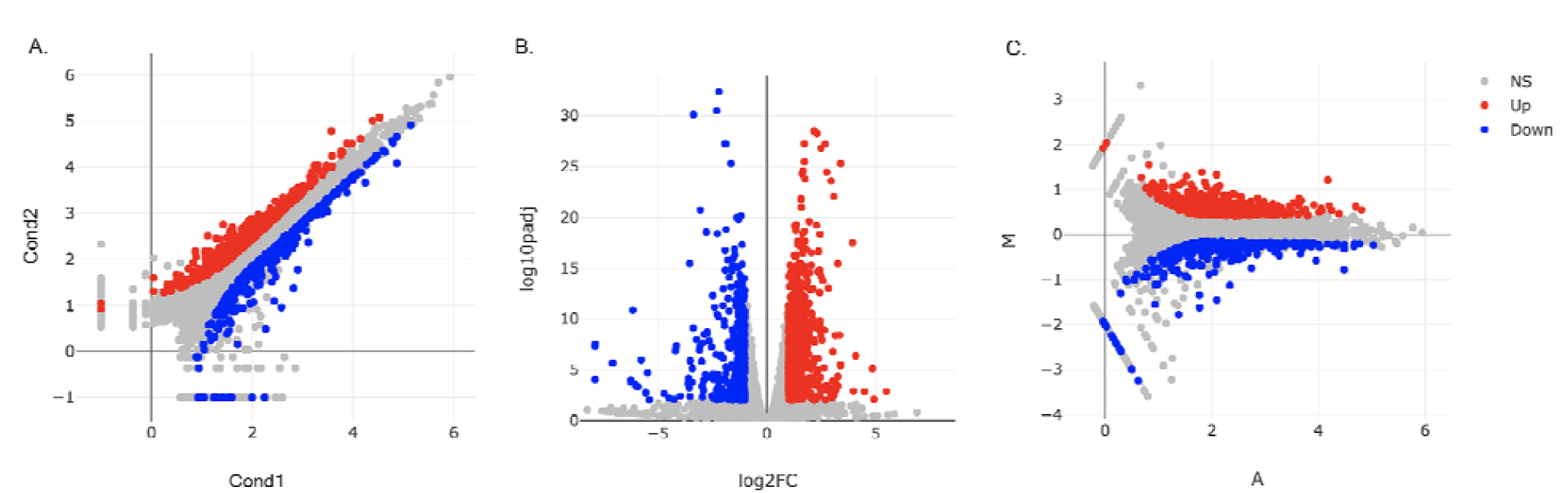
**a** Scatter plot. **b** Volcano plot. **c** MA plot DE genes are located in each plot while padj < 0.01 and |log2foldChange|>1.

As reported by the authors, the L^Δ2^ has a stronger effect in high fat diet fed animals compared to that of the L^Δ1^ using this table. Additionally, exploring genes that are specifically affected by each KO suggests that individual JNK1 and JNK2 might have distinct functions in addition to their partially redundant functions. To examine genes that are up-regulated in the liver under high fat diet fed mice, we performed DE analysis between WT mice fed with a normal or high fat diet. In all, 493 genes are significantly down-regulated and 350 are upregulated in the livers of mice fed with a high-fat diet (p<0.01, fold change > 1.5). Disease ontology analysis of upregulated genes shows, not surprisingly, a clear enrichment of diseases resulting from poor diet (urinary, kidney and other obesity related ailments). Moreover, enriched insulin signaling pathway genes are shown on a scatter plot (Fig. 6a) and a heatmap is created (Fig. 6b) using selected genes on this scatter plot. Additionally, differentially expressed genes from mice fed a chow diet and HFD (log2 fold ratio ≥ 0.75; p < 0.001) were examined by k-means (k = 6) clustering (Fig. 6c) and heatmap of PPARα pathway (Fig. 6d) is created which is similar to that in supplementary (Fig. S2a, S2b) in the original report [2].

**Fig. 6.**
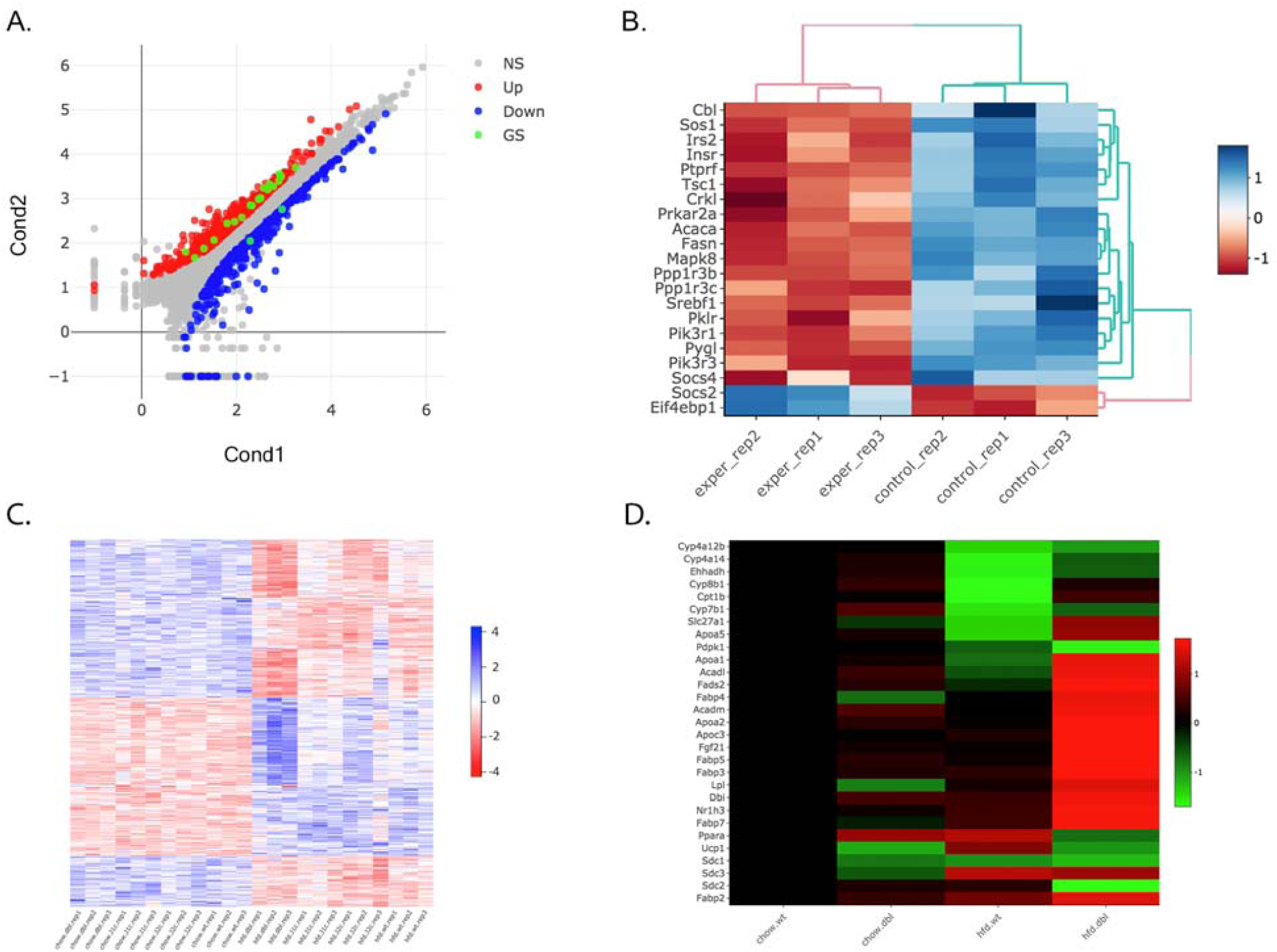
**a** Scatter plot of the genes enriched in insulin signaling pathway are located by green dots and **b** their expression values visualize in a heatmap. **c** Heatmap of differentially expressed genes from mice fed a chow diet and HFD (log2 fold ratio ≥ 0.75; p < 0.001) were examined by k-means (k = 6) clustering **d** PPARα pathway is presented as a heatmap where scale represents log2 fold change compared with chow-fed L^WT^ mice.

Further, users can explore individual regions by hovering over points. The gene or region id is shown and a barplot displaying the values of the gene or region across all samples is drawn. This is especially useful to investigate, for example, why certain genes may have large differences in between samples but fail to achieve statistical significance. In Fig. 7a, the, Fabp3 gene fails to achieve a significance of 0.05 despite its mean expression being between the conditions exceeding a 50 fold change. Hovering over Fabp3 shows the high variance of this gene across samples, which explains why statistical tests that account for both inter and intra condition variability to achieve significance in cases of high intra condition variability (Fig. 7b).

**Fig 7.**
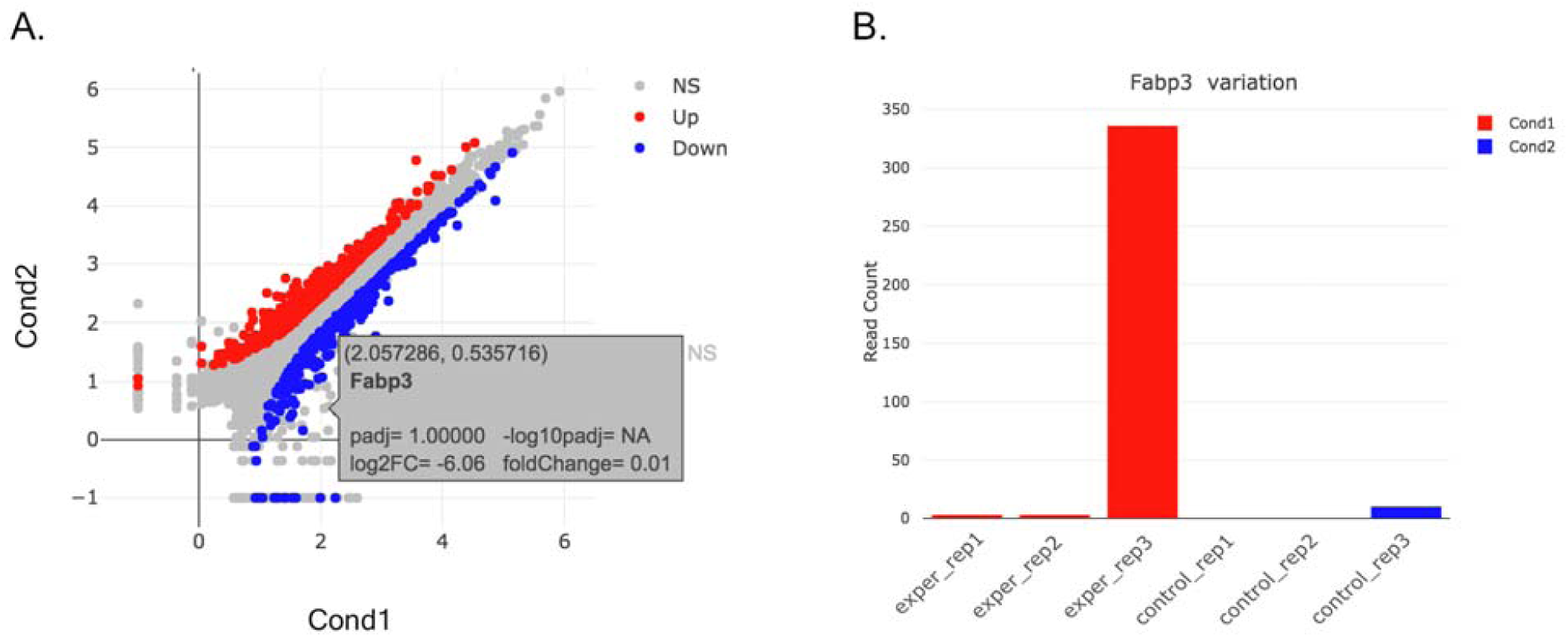
**a** FABP3 gene is hovered over on a scatter plot. **b** The read counts for FABP3 gene is shown as a bar plot.

Volcano, Scatter and MA plots work on summary statistics: significance, averages or fold change of averages. To explore the underlying data for any set of regions in a plot, DEBrowser can draw heatmaps for any selected region from any main plot. Selection can be made in a rectangular form or as a free-form using plotly’s lasso select (Fig. 8a). Fig. 8b shows the heatmap for the selected genes. Conversely, in any heatmap the users can select a subset of regions (such as based on similar expression pattern) for downstream analysis such as gene ontology, disease and pathway analysis.

**Fig. 8.**
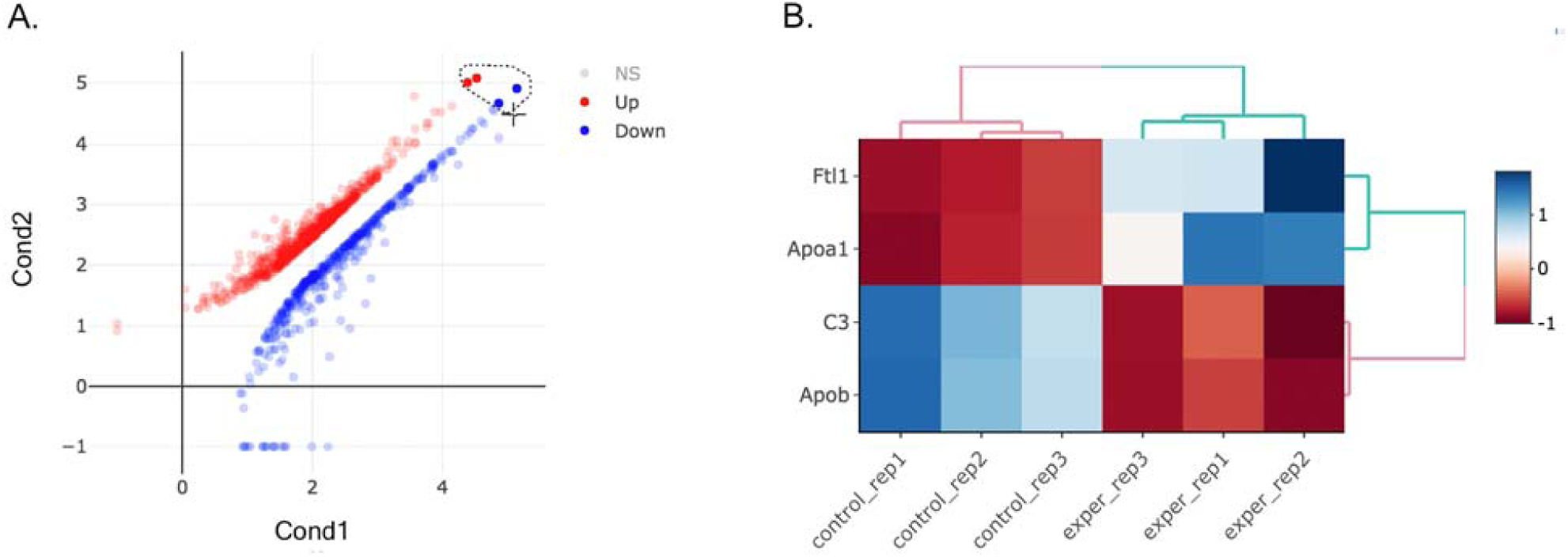
**a** Scatter plot of up and down regulated genes. **b** Heatmap of lasso selected of two up (Apob, C3) and two down regulated genes (Apoa1, Ftl1).

#### Gene ontology, disease and pathway discovery

For gene expression analysis in particular, DEBrowser supports Gene Ontology (GO) [53], KEGG pathway [53] and disease ontology analysis [54]. Users can perform GO or Pathway analysis directly on the results of differential expression analysis or on a subset of selected genes from any of the plots described above. For KEGG pathway analysis, in particular, DEBrowser provides pathway diagram for each enriched category (Fig. 9).

**Fig. 9.**
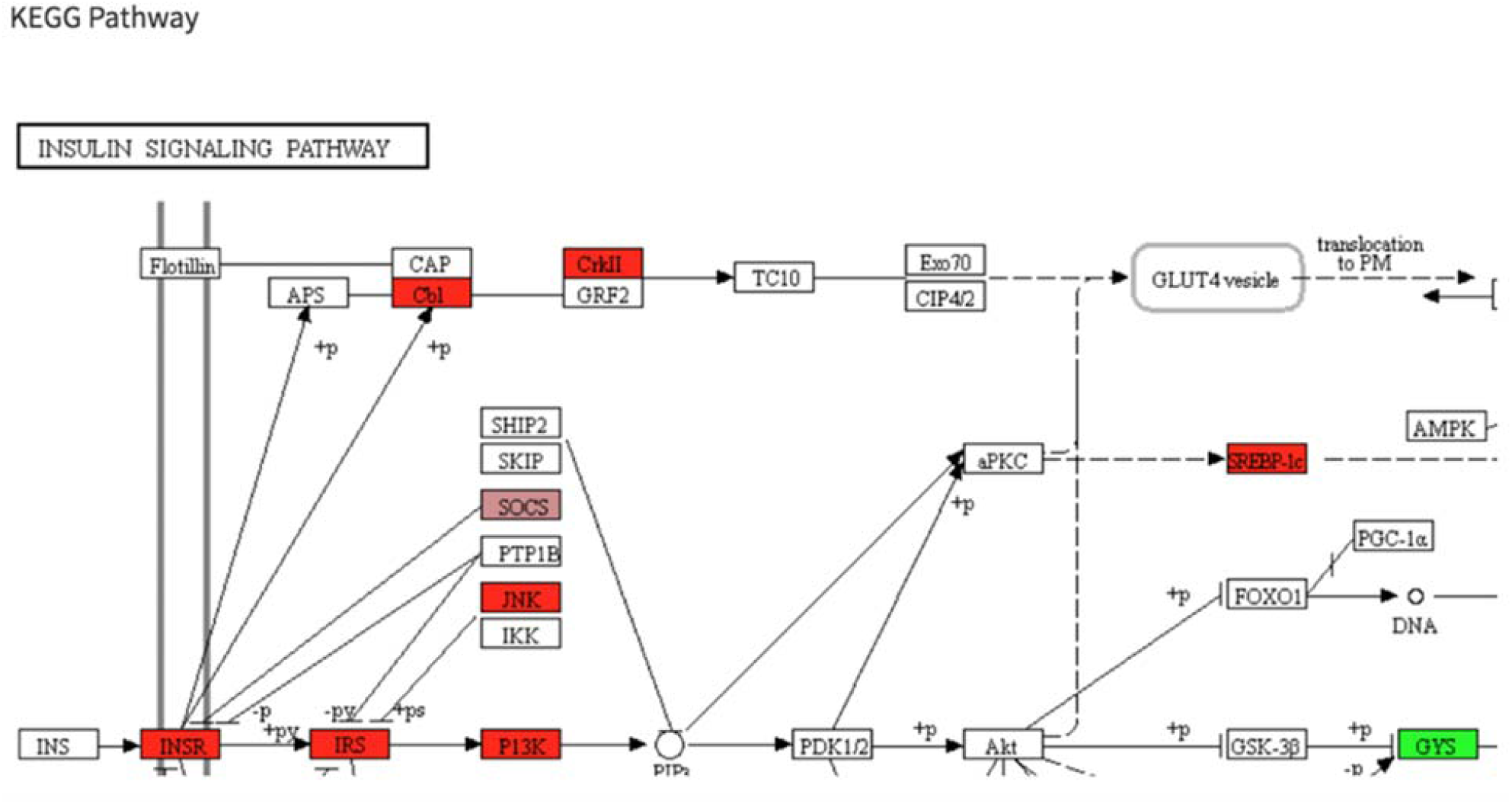
Insulin signalling pathway. Detected DE genes shown with colors according to their fold changes.

To further assist users in differential analysis, DEBrowser provides k-means clustering of differential regions, and when these regions are associated with genes, a gene ontology enrichment analysis is performed using enrichGO function in ClusterProfiler package [53] (Fig. 10).

**Fig. 10.**
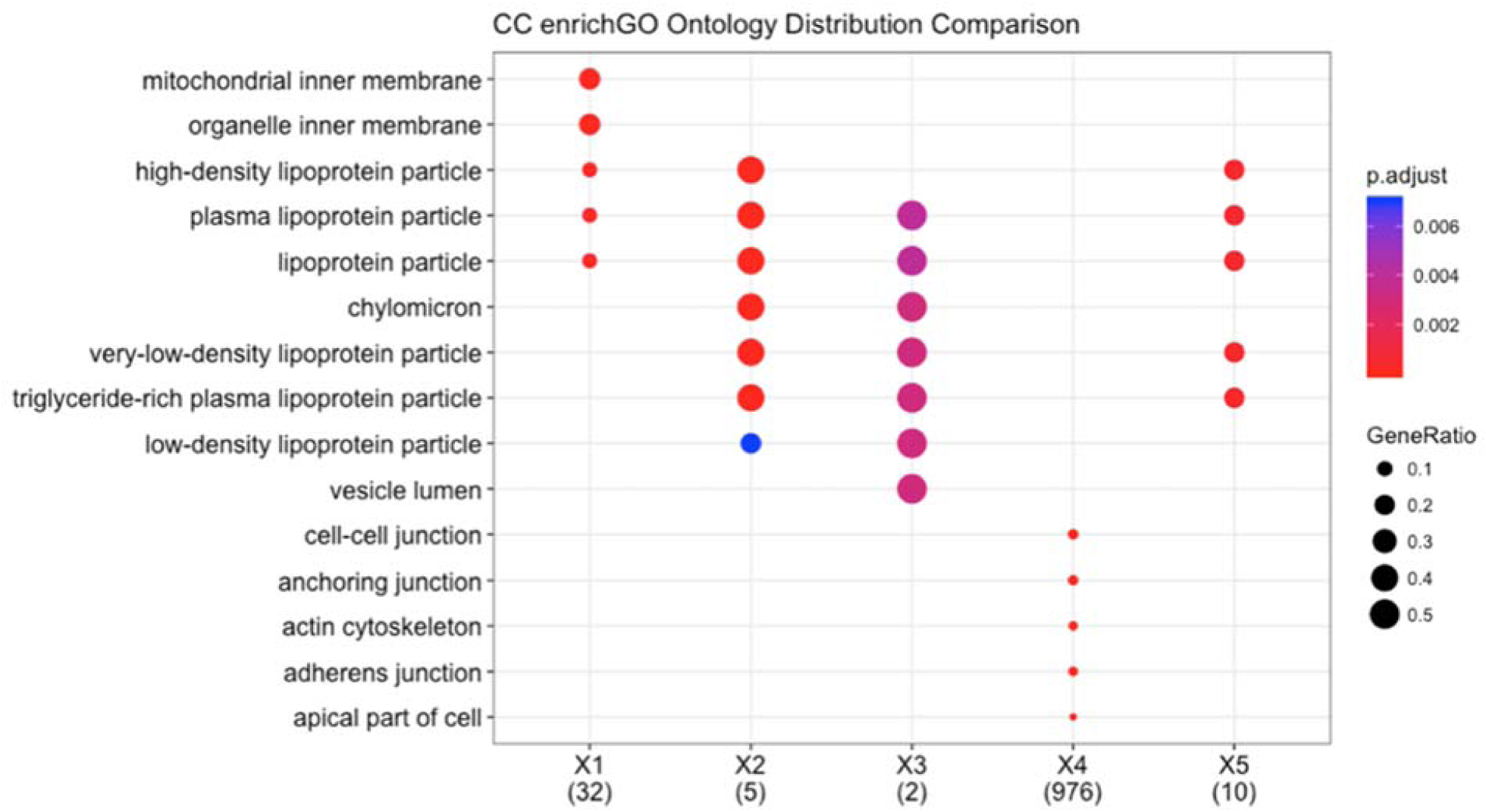
An example of a cluster profiler plot.

### Comparison to related applications

There are several applications, with varying functionalities, available for the exploration and analysis of DE. Most notable ones are, OASIS[27], VisRseq[28], DEGUST[29], DEIVA[30], WebMeV[31], Chipster[32], and DEapp[33]. A comparison of DEBrowser features to those applications is shown in Table S1.

### DEBrowser Modular Design

To reduce the code complexity and manage the program easier, while the DEBrowser features grow larger, we designed the components in a modular structure. To this end, bar, box and scatter plots, heatmaps with their corresponding controls can be reused multiple times in DEBrowser. We also shared sample applications that use individual modules. This modularity increased our development and test speed and code reusability. Such as, the size and the margins of the plots are controlled within the same module in all the plots in DEBrowser. This and other modules can be used in any other application that needs similar functionalities.

## Conclusion

The DEBrowser application provides users, who do not have any programming experience, with the ability to perform their own analysis. Therefore, it fills a void with a much needed graphical user interface for the analysis of count data that is typical of sequencing assays.

Existing tools do not include all of the steps in differential expression analysis and visualization. Another drawback of existing tools is that they usually assume that the data quality is good and is ready for differential expression analysis. Additionally, the plots are usually static and do not allow the interactivity necessary to understand the different parts of the data using different parameters real time, which reduces the efficiency of data exploration.

DEBrowser leverages open source components that are in active development in bioconductor [55,56], thus it benefits from a large community of developers. Its modular design makes it easy to swap components shall new paradigms or projects emerge that provide more ideal functionality than currently available.

## Declarations

### Acknowledgements

We would like to thank Kyle Jake Gellatly, Pranitha Vangala, Elisa Donnard, Rachel Murphy, Laney Zurlein, Stephen McGregor, Michael P. Czech, Leonardo Collado-Torres, and all members of the Garber Lab for their suggestions and comments.

### Funding

This work was supported by fund from the National Human Genome Research Institute NHGRI grant #U01 HG007910-01 and the National Center for Advancing Translational Sciences grant #UL1 TR001453-01 (M.G).

### Availability of data and materials

Table S1.docx: Application feature comparison table.

Additional file 1.pdf: Fig. S1 All2all scatter plot before (A) and after (B) batch correction

Additional file 2.pdf: Fig. S2 Heatmap of open chromatin regions affected by LPS stimulation.

Additional file 3.pdf: Installation instructions from bioconductors development repository.

### Authors’ contributions

AK implemented the package and drafted the manuscript. OY and DO interpreted data. MG supervised the project and critically revised the manuscript. All authors conceived the project, have read and approved the final manuscript.

### Ethics approval and consent to participate

Not applicable.

### Consent for publication

Not applicable.

### Competing interests

The authors declare that they have no competing interests.

## Abbreviations

RNA-Seq: RNA sequencing
ChIP-Seq: Chromatin immunoprecipitation sequencing
CLIP-Seq: Cross-linking immunoprecipitation sequencing
RIP-Seq: RNA immunoprecipitation sequencing
ATAC-Seq: Assay for transposase-accessible chromatin
PCA: Principal component analysis
hMDDCs: Human monocyte derived mouse dendritic cells
QC: Quality control
IQR: Interquartile range
MRN: Median ratio normalization
TMM: Trimmed Mean of M-values
RLE: Relative Log Expression
TLR: Toll-like receptor
JNK1 and JNK2: Jun terminal kinases
WT: Wild type
LΔ1: Jnk1 knockout
LΔ2: Jnk2 knockout
LΔ1,2: Jnk1 and Jnk2 double knockout
GO: Gene Ontology

## Availability and requirements

**Project name**: DEBrowser

**Project home page**: https://bioconductor.org/packages/release/bioc/html/debrowser.html, https://github.com/UMMS-Biocore/debrowser

**Documentation**: http://debrowser.readthedocs.org

**Operation system**s: Platform independent

**Programming language**: R (>□=□3.5.0)

**License**: GPL-v3

**Restrictions to use by non-academics**: None

